# A FOXA1/SPDEF Co-regulatory Axis Drives ERBB2 and Suppresses TEAD/YAP-Driven EMT to Maintain Luminal Identity in HER2+ Breast Cancer

**DOI:** 10.1101/2024.04.16.589460

**Authors:** Jaekwang Jeong, Jongwon Lee, Seungsoo Kim, Jaechul Lim, Jaehun Shin, Kwangmin Yoo, Jonghun Kim, Yoshiaki Tanaka, Hyun Seop Tae, Lark Kyun Kim, In-Hyun Park, Samir Zaidi, John Wysolmerski, Jungmin Choi

**Affiliations:** Section of Endocrinology and Metabolism, Department of Internal Medicine, Yale University School of Medicine, New Haven, CT, USA; Department of Biomedical Sciences, Korea University College of Medicine, Seoul, Republic of Korea; Brain Korea 21 Plus Project for Biomedical Science, Korea University College of Medicine, Seoul, Korea; College of Veterinary Medicine and Research Institute for Veterinary Science, Seoul National University, Seoul 08826, Republic of Korea; Integrated Science Engineering Division, Underwood International College, Yonsei University, Republic of Korea; Department of Genetics, Yale Stem Cell Center, Yale School of Medicine, New Haven, CT 06520, USA; Department of Medicine, Maisonneuve-Rosemont Hospital Research Centre, University of Montreal, Montreal, Quebec, H1T 2M4, Canada; Elgen Therapeutics, 193 Munji-Ro, Yuseong-Gu, Daejeon 34051, South Korea; Department of Biomedical Sciences, Graduate School of Medical Science, Brain Korea 21 Project, Gangnam Severance Hospital, Yonsei University College of Medicine, Seoul 06230, Republic of Korea; Department of Medicine, Division of Solid Tumor Oncology, Genitourinary Oncology Service, Memorial Sloan Kettering Cancer Center, New York, NY 10065, USA

**Keywords:** Breast Cancer, FOXA1, SPDEF, HER2, ERBB2, Epithelial-Mesenchymal Transition (EMT), TEAD/YAP, TRPS1

## Abstract

The function of the pioneer factor forkhead box A1 (FOXA1) in estrogen receptor (ER)-negative, human epidermal growth factor receptor 2 (HER2, encoded by the *ERBB2* gene)-positive breast cancer represents a critical gap in our understanding of lineage identity in tumorigenesis. Here, we solve this enigma by addressing this question through the identification of an indispensable, co-dependent regulatory circuit formed between FOXA1 and the SAM pointed domain containing ETS-family transcription factor (SPDEF). Using integrative multi-omics analyses of patient tumors and experimental models, we demonstrate that this circuit functions as the master guardian of the HER2-positive luminal state. It executes this role via two distinct, essential mechanisms: it directly binds regulatory elements to drive high-level expression of the *ERBB2* oncogene, and it simultaneously preserves epithelial identity by suppressing a latent program of epithelial-to-mesenchymal transition (EMT). Mechanistically, the FOXA1/SPDEF circuit promotes the expression of the transcriptional repressor TRPS1, which in turn antagonizes the pro-mesenchymal activity of the TEAD/YAP complex. Our findings define a central regulatory node that couples lineage fidelity to oncogenic output in HER2-positive cancer, revealing the FOXA1/SPDEF–TRPS1 axis as a critical determinant of tumor plasticity and a potential therapeutic target for overcoming resistance in HER2-positive breast cancer.

## Introduction

Forkhead box protein A1 (FOXA1) and SAM pointed domain-containing ETS transcriptional factor (SPDEF) are luminal-lineage transcription factors that play central roles in establishing and maintaining epithelial identity in hormone-responsive tissues, including the mammary gland and prostate (1–3). FOXA1 is widely recognized as a pioneer factor that binds condensed chromatin and generates accessible regulatory regions, thereby licensing transcriptional programs associated with luminal differentiation (4–6). In ER-positive breast tumors, this function is inextricably linked to the estrogen receptor (ERα, encoded by the *ESR1* gene); FOXA1 facilitates ERα’s access to gene promoters, anchoring the estrogen-responsive transcriptional programs that define the canonical luminal phenotype (4, 7, 8). However, a significant subset of human epidermal growth factor receptor 2 (HER2, encoded by the *ERBB2* gene)-positive tumors is classified as luminal and expresses high levels of FOXA1 yet lacks ERα(9, 10). This presents a critical enigma in breast cancer biology: what is the primary function of FOXA1 in the absence of its canonical partner, and what molecular machinery does it employ to establish and defend the luminal state?

SPDEF is an ETS family transcription factor involved in differentiation and maturation in epithelial tissues such as the intestine, prostate, and mammary gland (11–13). It has been reported that FOXA1 directly regulates *SPDEF* gene expression in several tissues (14–16). Within this context, SPDEF is recruited to a subset of FOXA1-primed enhancers, where it reinforces epithelial gene expression and promotes terminal differentiation programs such as secretion and maintenance of epithelial polarity (14, 15). Given its established role as a downstream effector of FOXA1 and its complex, context-dependent functions in breast cancer—acting as both a tumor promoter and suppressor (15, 17, 18) —SPDEF represents a logical candidate to cooperate with FOXA1 in an ER-independent context.

HER2 amplification is another major driver of breast cancer and defines a distinct clinical subtype (19, 20). The output of HER2 signaling is strongly influenced by transcription factors. Luminal transcription factors such as FOXA1 and SPDEF contribute to the maintenance of epithelial identity and create a chromatin environment that supports receptor tyrosine kinase signaling. Although FOXA1 has been shown to regulate *ERBB2* transcription directly, the precise mechanisms and cofactors involved remain unclear (21). Here, we test the hypothesis that FOXA1 and SPDEF form a co-dependent, ER-independent regulatory axis that functions as the master guardian of the HER2-positive luminal state. We propose that this axis is essential for both directly maintaining the expression of the lineage-defining oncogene *ERBB2* and actively suppressing a latent program of EMT to preserve luminal lineage fidelity.

## Results

### Reciprocal co-dependency of FOXA1 and SPDEF in HER2-positive breast cancer

FOXA1 is well known to promote ERα gene expression and is necessary for the ER-responsive transcriptome of canonical ER-positive luminal breast cancers (4, 22). However, FOXA1 is also expressed in HER2-positive breast cancers with low levels of ERα. To assess the effects of knocking out *FOXA1*, we examined a comprehensive panel of cell lines (n = 1,225) from the Cancer Dependency Map (23, 24). This analysis revealed that HER2-positive breast cancer cells rely on FOXA1 for viability, indicated by aggregate gene effect scores around -1 in the absence of *FOXA1* (a score of –1 is generally considered strongly lethal; *SI Appendix*, Fig. S1A). Because correlations between dependency profiles often indicate shared pathway functionality, we evaluated genes co-dependent with *FOXA1*. *SPDEF* emerged as the gene most strongly correlated with *FOXA1* dependency, and vice versa (Fig. 1A). In fact, among all genes, *FOXA1* and *SPDEF* the highest co-dependency correlation specifically in HER2-positive breast cancer.

**Figure 1.**
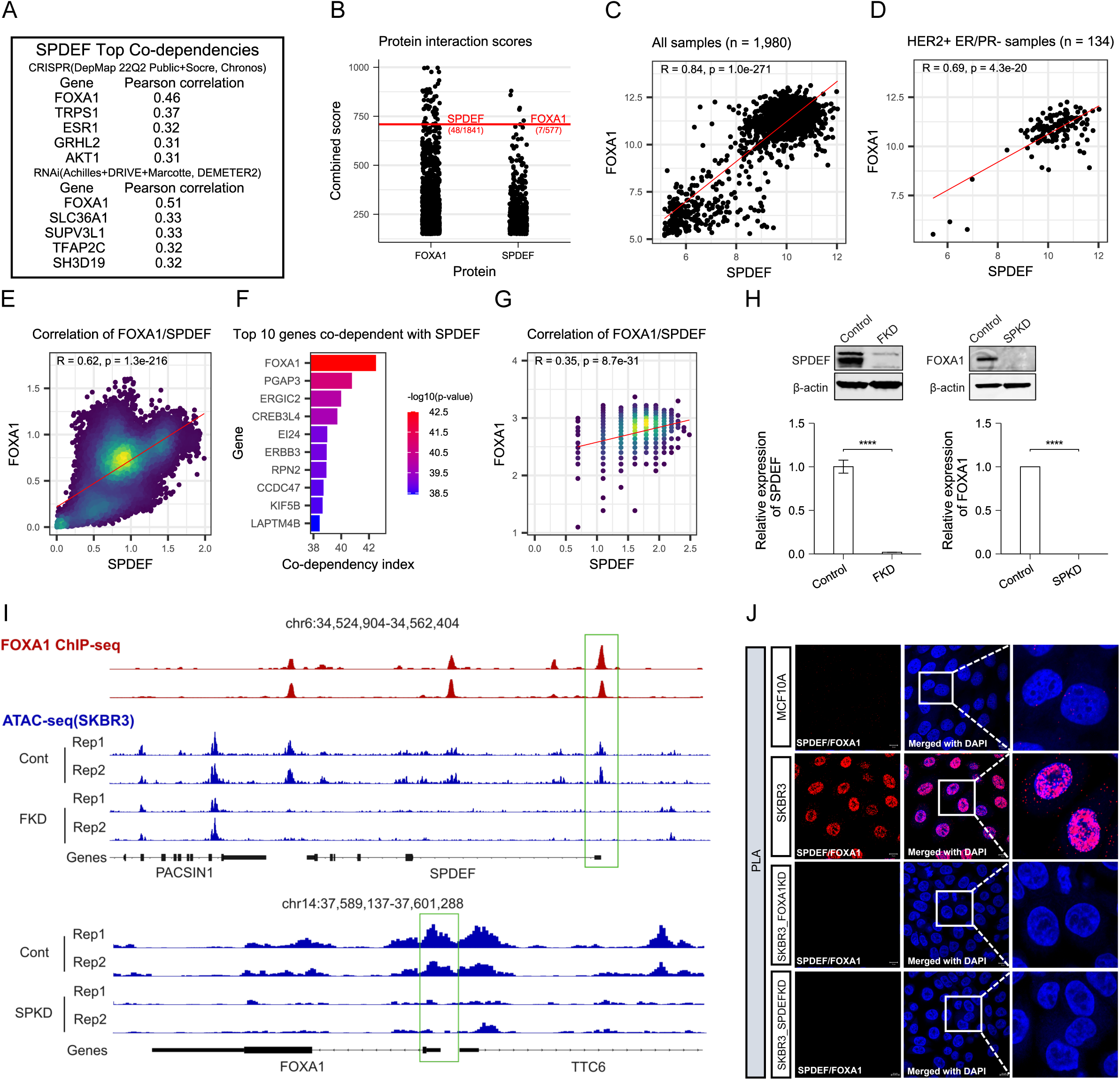
Reciprocal Co-dependency of FOXA1 and SPDEF in HER2-positive breast cancer. **A**. Top SPDEF correlated genes from RNAi and CRISPR screening data for 1,255 cell lines. **B**. Jittered plot showing the interaction scores between FOXA1 and SPDEF derived from a protein interaction database. **C-D**. Scatter plot depicting FOXA1 and SPDEF gene expression level in the entire cohort or HER2+ ER/PR- of breast cancer patients from the METABRIC dataset (C, n = 1,980 and D, n = 134, respectively). **E**. Scatter plot showing the correlation between FOXA1 and SPDEF expression in breast cancer scRNA-seq data. **F**. Bar plot displaying the top 10 genes showing co-dependency with SPDEF based on scRNA-seq data. **G**. Scatter plot showing the correlation between FOXA1 and SPDEF expression in Visium spatial transcriptomic data. **H**. Western blots and histograms showing the expression levels of SPDEF and FOXA1 in the knockdown cell line. **I**. Genome tracks of SPDEF and FOXA1 gene promoter region showing the difference in accessibility between control and knockdown cells. FOXA1 binding motifs matched with ChIP-seq peak regions were marked with a green square. **J**. Proximity Ligation Assay (PLA) assay of SPDEF/FOXA1 in MCF10A, SKBR3, SKBR3-FKD and SKBR3-SPKD cells. Nuclei stained with DAPI (blue). Scale bars are 10μm. Bar graphs represent the mean±SEM. **** denotes p<0.00005.

To investigate the association between FOXA1 and SPDEF in HER2-positive breast cancer, we first examined protein interaction data. FOXA1 and SPDEF exhibited a strong interaction (interaction scores 48/1841 and 7/577, respectively, in a protein-interaction database), indicating a significant relationship (Fig. 1B). These scores reflect a notable frequency of reported FOXA1-SPDEF interactions, suggesting potential cooperation of FOXA1 and SPDEF within regulatory networks relevant to breast cancer biology. To validate this association at the transcriptomic level, we analyzed the METABRIC dataset, one of the largest breast cancer cohorts. Both the overall patient cohort (Fig. 1C) and the HER2-positive ER/Progesterone receptor (PR)-negative subgroup (Fig. 1D) showed a high degree of correlation between *FOXA1* and *SPDEF* expression, with Pearson correlation coefficients of R = 0.84 (p = 1.0 × 10^-271^) and R = 0.69 (p = 4.3 × 10^-20^), respectively. These findings highlight a robust positive coupling of *FOXA1* and *SPDEF* expression, consistent with a cooperative role in HER2-positive breast cancer.

Next, we validated these findings using single-cell RNA sequencing (scRNA-seq) and spatial transcriptomics. Reanalysis of a published scRNA-seq dataset from 19 human primary breast tumors, comprising 6 HER2-positive and 13 ER-positive (25) revealed three major cell populations: epithelial, immune, and stromal (*SI Appendix*, Fig. S1B–D). *FOXA1* and *SPDEF* were strongly co-expressed in the epithelial compartment; we observed a Pearson correlation of R = 0.62 (p = 1.3 × 10^-216^) between *FOXA1* and *SPDEF* transcripts at the single-cell level (Fig. 1E). Co-dependency analysis of these single-cell data again identified *FOXA1* as the top-ranking gene co-dependent with *SPDEF* (Fig. 1F).

To elucidate the spatial co-expression patterns of *ERBB2*, *FOXA1*, and *SPDEF* in tumors, we examined a 10X Visium spatial transcriptomics dataset of a human breast cancer specimen (ER-positive/PR-negative/HER2-positive, Stage IIB FFPE section) (26). A subsequent comparison of gene expression between the control group and the two tumor subtypes revealed higher co-expression density for *FOXA1* and *SPDEF*, aligning with the patterns seen in the *SI Appendix*, Fig. S1E and F. Notably, we observed co-localization of *FOXA1* and *SPDEF* expression in tumor regions (*SI Appendix*, Fig. S1E and F). Across all spatial spots, *FOXA1* and *SPDEF* showed a positive correlation [R = 0.35 (Pearson), p-value = 8.7 × 10^-31^], mirroring the single-cell results (Fig 1G). These scRNA-seq and spatial analyses support a close *FOXA1*–*SPDEF* association in HER2-positive breast cancer biology.

To investigate the reciprocal influence between FOXA1 and SPDEF, we generated stable knockdown models for each gene in the HER2-positive cell line SKBR3 (hereafter referred to as SKBR3-FKD and SKBR3-SPKD, respectively). Knockdown of *FOXA1* led to a significant reduction in *SPDEF* expression, and conversely, *SPDEF* knockdown reduced *FOXA1* levels (Fig. 1H), suggesting a regulatory interdependence. We then asked whether these changes in transcript/protein were linked to altered chromatin accessibility at the corresponding gene loci. ATAC-seq analysis revealed that *FOXA1* knockdown caused decreased chromatin accessibility at the *SPDEF* promoter. Similarly, *SPDEF* knockdown reduced accessibility at the *FOXA1* promoter (Fig. 1I). Finally, we tested if FOXA1 and SPDEF physically interact. Using a proximity ligation assay (PLA) for FOXA1–SPDEF interactions, we found abundant nuclear PLA signals in control SKBR3 cells, which were markedly lost upon knockdown of either *FOXA1* or *SPDEF* (Fig. 1J). Together, these findings provide compelling evidence of a co-dependent and reciprocal regulatory relationship between FOXA1 and SPDEF in HER2-positive breast cancer, establishing the existence of a core FOXA1/SPDEF circuit.

### FOXA1 and SPDEF are required for *ERBB2* expression in HER2-positive breast cancer cells

We next examined whether FOXA1 and SPDEF affect *ERBB2* expression and the downstream HER2 signaling pathway. Using the SKBR3-FKD and SKBR3-SPKD models, we found that silencing either *FOXA1* or *SPDEF* markedly reduced *ERBB2* mRNA levels (by quantitative RT-PCR) and HER2 protein levels (by immunoblot) relative to control (Fig. 2A and B). Correspondingly, the level of phosphorylated HER2 (pHER2) and phosphorylated epidermal growth factor receptor (pEGFR) was significantly diminished (Fig. 2B). These findings suggest that FOXA1 and SPDEF are required to play a role in regulating HER2 expression and signaling in HER2-positive cells.

**Figure 2.**
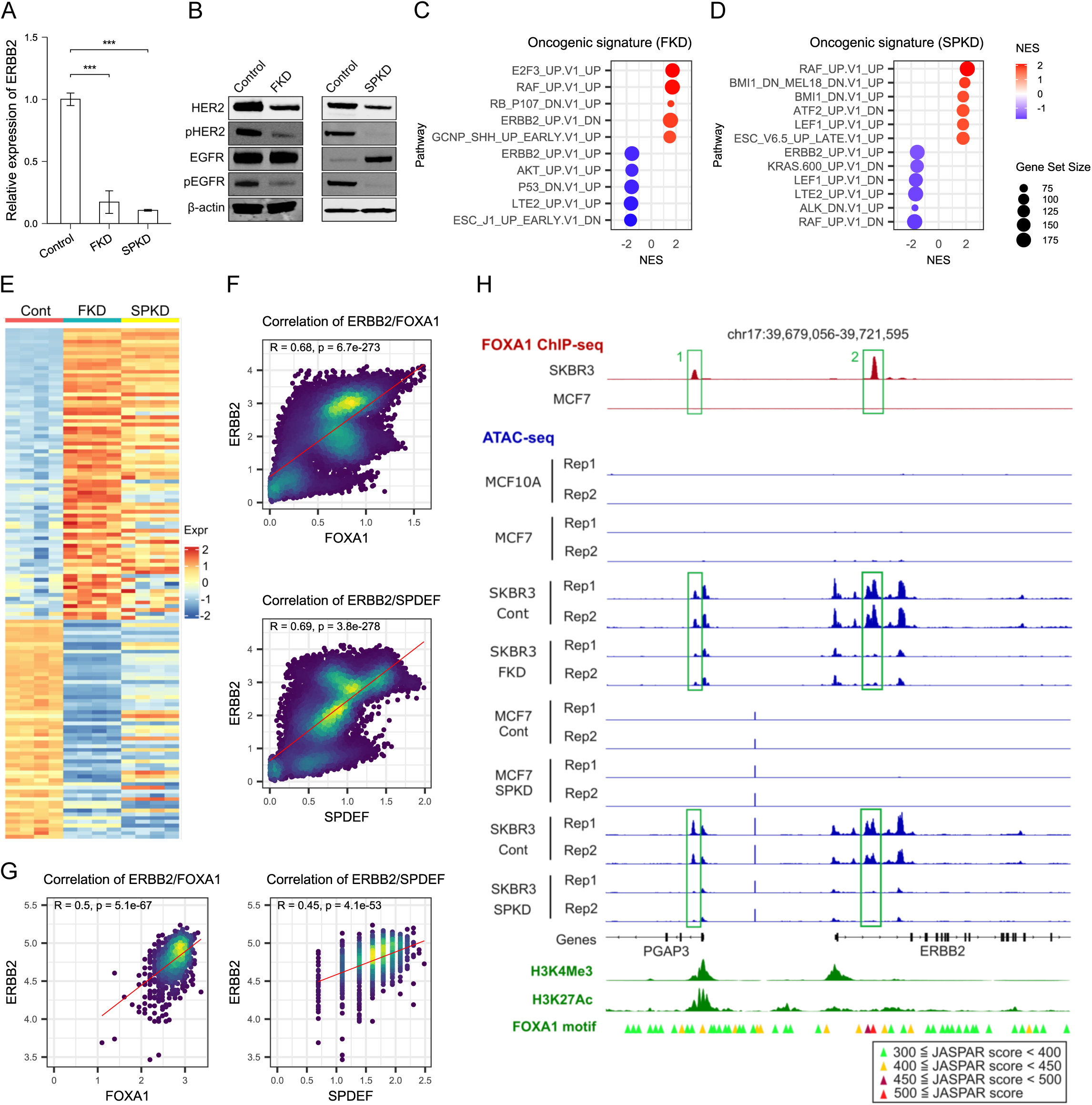
FOXA1 and SPDEF are required for *ERBB2* expression in HER2-positive breast cancer cells. **A**. Histogram showing the relative expression levels of *ERBB2* from qRT-PCR in SKBR3, SKBR3-FKD, and SKBR3-SPKD cells. **B**. Western blots showing the indicated protein levels in SKBR3, SKBR3-FKD, and SKBR3-SPKD cells. **C**. Dot plot illustrating the top 10 enriched terms of GSEA in the analysis of SKBR3-FKD cells compared to the SKBR3. **D**. Dot plot illustrating the top 12 enriched terms in the analysis of SKBR3-SPKD compared to the SKBR3. **E**. Heatmap showing the relative expression of ERBB2_UP.V1_DN leading-edge subset genes and ERBB2_UP.V1_UP leading-edge subset genes in SKBR3, SKBR3-FKD, and SKBR3-SPKD cells. **F**. Scatter plots showing the Pearson correlation coefficient between *FOXA1* and *ERBB2* among scRNA-seq data (upper panel). Between *SPDEF* and *ERBB2* (lower panel). P-value was calculated from a down-sampled dataset with 1 to 10 % to avoid zero inflation. **G**. Scatter plots showing the Pearson correlation coefficient between *FOXA1* and *ERBB2* among Visium data (left panel). Between *SPDEF* and *ERBB2* (right panel). **H**. Genome tracks of *ERBB2* gene promoter region showing the difference accessibility between control and knockdown cells. Genome tracks showing the identified peaks over pre-defined promoter region (H3K4Me3) and regulatory region (H3K27Ac). FOXA1 binding motifs matched with ChIP-seq peak regions were marked with a green square. Triangles colored according to JASPAR binding affinity scores mark the locations of FOXA1 binding motifs within the promoter site. Bar graphs represent the mean±SEM. *** denotes p<0.0005, **** denotes p<0.00005.

To gauge the global effects of FOXA1/SPDEF loss on HER2 signaling pathway, we performed Gene Set Enrichment Analysis (GSEA) on RNA-seq data (*SI Appendix*, Dataset S1 and S2). Oncogenic signature GSEA revealed that HER2 signaling outputs were among the top five gene sets downregulated in both SKBR3-FKD and SKBR3-SPKD cells (Fig. 2C and D; *SI Appendix*, Dataset S3 and S4). Specifically, the “ERBB2_UP.V1_UP” gene set (190 genes upregulated by constitutively active HER2) was significantly negatively enriched (normalized enrichment score [NES] = -1.57, adjusted p-value = 1.59 × 10^-3^ for SKBR3-FKD; NES = -1.56, adjusted p-value = 2.06 × 10^-3^ for SKBR3-SPKD). Similarly, the “AKT_UP.V1_UP” gene set (NES = -1.57, adjusted p-value = 1.6 × 10^-3^ for SKBR3-FKD; NES = -1.51, adjusted p-value = 7.95 × 10^-3^ for SKBR3-SPKD) was negatively enriched, consistent with suppressed cell growth and survival signaling upon *FOXA1* and *SPDEF* loss. Conversely, the “ERBB2_UP.V1_DN” gene set (200 genes normally repressed by constitutive active HER2) was positively enriched in SKBR3-FKD cells (NES = 1.55, adjusted p-value = 2.7 × 10^-3^) (Fig. 2C). An orthogonal over-representation analysis (ORA) of differentially expressed genes yielded similar conclusions (*SI Appendix*, Fig. S2A and B, Dataset S5 and S6). A heatmap of leading-edge genes from the HER2 signature gene sets highlights the definitive role of FOXA1 and SPDEF in regulating HER2-related transcriptional programs (Fig. 2E). Key genes from these signatures showed reciprocal changes (up or down) consistent with diminished HER2 signaling (*SI Appendix*, Fig. S2C and D).

We also examined *FOXA1*/*SPDEF* association with *ERBB2* expression in the patient datasets. In scRNA-seq data, *FOXA1* and *ERBB2* transcripts were strongly positively correlated [R = 0.68 (Pearson), p = 6.7 × 10^-273^], as were *SPDEF* and *ERBB2* [R = 0.69 (Pearson), p = 3.8 × 10^-278^] (Fig. 2F). Spatial transcriptomics analysis similarly revealed that the spots with high *FOXA1* or *SPDEF* tended to have high *ERBB2* [R = 0.5 (Pearson), p = 5.1 × 10^-67^ for *FOXA1*; R = 0.45 (Pearson), p = 4.1 × 10^-53^ for *SPDEF*] (Fig. 2G). These *ex vivo* observations align with our experimental findings, which show that FOXA1 and SPDEF sustain *ERBB2* expression.

To determine whether FOXA1 and SPDEF directly regulate *ERBB2*, we performed FOXA1 ChIP-seq in SKBR3 cells. We identified two distinct, narrow FOXA1 binding peaks in the *ERBB2* locus. Notably, these peaks were not in the canonical proximal promoter region of *ERBB2*. One peak (labelled “1” in Fig. 2H) was located in a region that corresponds to either the *ERBB2* or an adjacent *PGAP3* promoter (according to the eukaryotic promoter database), and the second peak (“2”) lay downstream of transcription start site (TSS) of the canonical *ERBB2* transcript, defining a putative enhancer (Fig. 2H). Importantly, ATAC-seq data showed that these same two regions were in open chromatin in SKBR3 cells, overlapping the FOXA1 ChIP-seq peaks. This suggests that FOXA1 binds to accessible regulatory DNA within *ERBB2*, likely driving its expression. Consistently, we observed that these *ERBB2* promoter/enhancer peaks lost accessibility in SKBR3-FKD or SKBR3-SPKD cells (Fig. 2H). By contrast, in ER-positive MCF7 cells (which express FOXA1 but not HER2), no FOXA1 binding was detected at *ERBB2*. Instead, FOXA1 bound the *ESR1* gene promoter (estrogen receptor) in MCF7 (*SI Appendix*, Fig. S2E), highlighting a context-specific targeting: FOXA1 binds *ESR1* gene regulatory regions in ER-positive cells but *ERBB2* regions in HER2-driven cells (Fig. 2H). No *ERBB2* promoter binding or accessible chromatin was seen in HER2-negative, non-transformed human mammary epithelial cell lines MCF10A or MCF7 (Fig. 2H). Collectively, these findings provide compelling evidence that FOXA1 and SPDEF directly regulate *ERBB2* transcription by modulating its promoter/enhancer accessibility, thereby controlling HER2 expression and downstream signaling.

### FOXA1 and SPDEF are required for luminal identity in HER2-positive breast cancer

FOXA1 and SPDEF contribute to maintaining the luminal epithelial phenotype, thus, loss of FOXA1 and SPDEF might promote a shift toward basal-like characteristics in HER2-positive tumors. To test this hypothesis, we identified differentially expressed genes (DEGs) from RNA-seq of SKBR3-FKD and SKBR3-SPKD compared to parental SKBR3. Among the top five upregulated genes in SKBR3-FKD (*SI Appendix*, Dataset S1), four (*ABCG2*, *COL14A1*, *FSCN1*, and *SLC2A3*) have previously been associated with drug resistance, disease progression, metastasis, and EMT in breast cancer (27–30). Similarly, of the top five genes induced by SKBR3-SPKD (*SI Appendix*, Dataset S2), three (*SPARC*, *HTRA1*, and *ADAMTS1*) have been reported to play roles in extracellular matrix turnover, cancer cell proliferation, invasion, and EMT (31–33).

To assess the global transcriptomic shifts, we performed ORA on the DEGs and GSEA. Using a stringent cutoff (|log2 fold change| > 3), ORA showed that Hallmark gene sets related to EMT, angiogenesis, and inflammatory response were significantly enriched among gene upregulated in both SKBR3-FKD and SKBR3-SPKD cells (Fig. 3A and B; *SI Appendix*, Dataset S7 and S8). When considering all DEGs without fold-change cutoff, we found that SKBR3-FKD cells upregulated genes involved in proliferation gene sets (MYC target, E2F target, G2M checkpoint) and WNT/β-catenin signaling (*SI Appendix*, Fig. S3A and C, Dataset S9 and S11), whereas SKBR3-SPKD cells upregulated genes in angiogenesis, EMT, and interferon-α response (*SI Appendix*, Fig. S3B and D, Dataset S10 and S12). Focusing on the EMT program specifically, genes from the Hallmark EMT set were broadly induced by loss of FOXA1/SPDEF. A heatmap of the leading-edge genes from the “HALLMARK_EMT” gene set and volcano plots (Fig. 3C; *SI Appendix*, Fig. S3E and F) illustrate that many epithelial markers (*MUC1*, *KRT4* and *KRT19*, etc.) were down, and mesenchymal markers (*VIM*, *CDH2* and *SERPINE2*, etc.) were up in both SKBR3-FKD and SKBR3-SPKD cells. These observations indicate that FOXA1 and SPDEF help sustain luminal characteristics, at least in part, by repressing EMT-associated genes in HER2-positive breast cancer.

**Figure 3.**
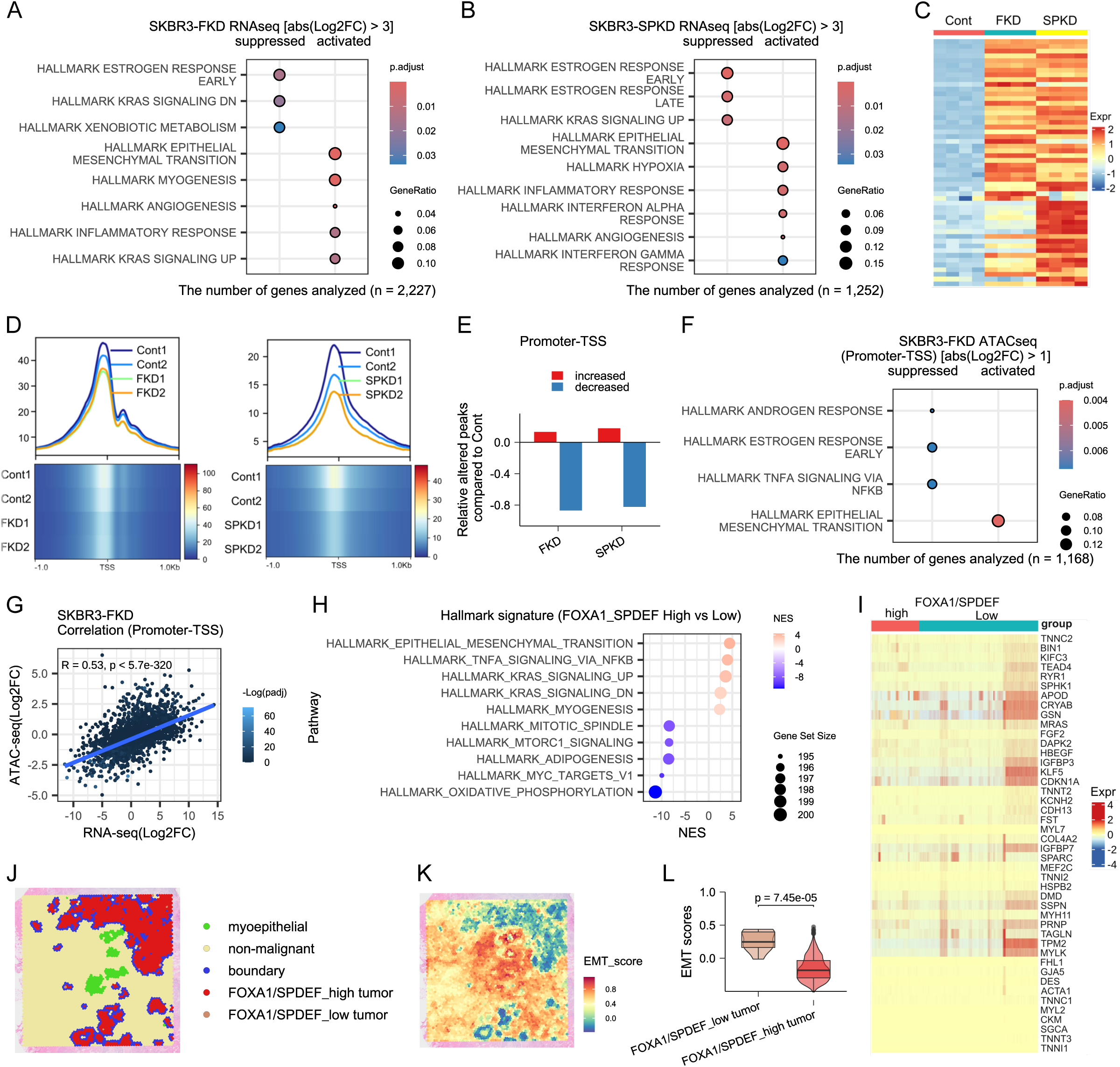
FOXA1 and SPDEF are required for luminal identity in HER2-positive breast cancer. **A-B**. Dot plot showing the significantly enriched Hallmark gene sets by ORA performed for DEGs (|Log2FC| > 3) in the SKBR3-FKD cells (A) or SKBR3-SPKD (B) compared to the SKBR3. **C**. Heatmap showing the relative expression of HALLMARK_EMT leading-edge subset genes in SKBR3, SKBR3-FKD, and SKBR3-SPKD cells. **D**. Tornado plot illustrating the differential chromatin accessibility peaks identified by ATAC-seq between control and knockdown conditions. ATAC-seq reads in 2-kb regions centered on the TSS sites of annotated promoter-TSS sites of protein-coding genes. **E**. Bar plot showing regions of chromatin accessibility altering between conditions. **F**. Dot plot showing the significantly enriched Hallmark gene sets by ORA performed for DEGs (|Log2FC| > 1) in the SKBR3-FKD cells compared to the SKBR3. Genes corresponded with DARs annotated only as Promoter-TSS were used for analysis. **G**. Scatter plot showing the correlation of the log2FC values between RNA-seq and ATAC-seq of SKBR3-FKD. Displayed R value determined by Pearson correlation. **H**. Dot plot showing the significantly enriched Hallmark gene sets by GSEA between *FOXA1*/*SPDEF*^low^ and *FOXA1*/*SPDEF*^high^ cells. **I**. Heatmap showing the relative expression of HALLMARK_EMT leading-edge subset genes between *FOXA1*/*SPDEF*^low^ and *FOXA1*/*SPDEF*^high^ cells. **J**. Spatial distribution of annotated cell types detected in Visium spatial transcriptomics data. **K**. Visualization of EMT gene module scores across the Visium spatial transcriptomics data. **L**. Violin plot illustrating the distribution of EMT scores in *FOXA1*/*SPDEF*^high^ and *FOXA1*/*SPDEF*^low^ expression groups. The box centerlines depict the medians, and the edges depict the first/third quartiles.

We next examined whether FOXA1/SPDEF loss also alters chromatin accessibility in line with an EMT shift. ATAC-seq comparing SKBR3-FKD and SKBR3-SPKD cells to control revealed widespread changes. Both *FOXA1* and *SPDEF* knockdown led to a global reduction in chromatin accessibility relative to the parental SKBR3, especially at promoter regions (Fig. 3D; *SI Appendix*, Fig. S3G and H, Dataset S13 and S14). While most promoter-TSS regions showed decreased accessibility, a subset of loci gained accessibility (Fig. 3E), suggesting selective chromatin opening at genes potentially linked to altered transcriptional programs. We next investigated the functional pathways influenced by these chromatin accessibility changes via ORA and GSEA. ORA of genes associated with differentially accessible regions (DARs) identified significant enrichment of the Hallmark EMT gene set in both SKBR3-FKD and SKBR3-SPKD (Fig. 3F; *SI Appendix*, Fig. S3I, Dataset S15 and S16). This robust enrichment of an EMT signature persisted when the analysis was expanded to include all genes associated with DARs (*SI Appendix*, Fig. S3J and K, Dataset S17 and S18), strongly implicating the activation of an EMT program as a central consequence of the chromatin accessibility changes.

To directly link DARs and gene expression, we correlated promoter accessibility with gene expression changes, using a previously reported method (34). In SKBR3-FKD, across the whole transcriptome, promoter accessibility correlated positively with gene expression [R = 0.53 (Pearson), P < 5.7 × 10^-320^] (Fig. 3G). A similar correlation, though slightly weaker, was observed in SKBR3-SPKD cells [R = 0.43 (Pearson), P < 3 × 10^-200^] (*SI Appendix*, Fig. S3L). Strikingly, when we restricted this analysis to a set of 73 Hallmark EMT-related genes that showed promoter accessibility changes, the correlation between chromatin opening and gene upregulation became even stronger for SKBR3-FKD cells [R = 0.63 (Pearson), P = 5.6 × 10^-9^] and SKBR3-SPKD cells [R = 0.671 (Pearson), P = 1 × 10^-9^] (*SI Appendix*, Fig. S3M and N), indicating genes driving EMT tended to both gain promoter accessibility and increase expression when FOXA1/SPDEF were lost.

Given that clinical tumors are heterogeneous, we asked if cells with low *FOXA1*/*SPDEF* within tumors show signs of EMT activation. We divided the epithelial tumor cells in the scRNA-seq data into *FOXA1*/*SPDEF*^high^ and *FOXA1*/*SPDEF*^low^ groups (see Methods). The *FOXA1*/*SPDEF*^low^ cells were indeed enriched for EMT-related genes (NES = 4.43, adjusted p-value = 1.38 × 10^-3^; Fig. 3H; *SI Appendix*, Dataset S19). A heatmap of the leading-edge genes from the “HALLMARK_EMT” gene set confirms that *FOXA1*/*SPDEF*^low^ population in patient samples expresses higher levels of mesenchymal markers (Fig. 3I). Consistently, in the spatial transcriptomics data, tumor spots with high *FOXA1*/*SPDEF* expression exhibited significantly lower EMT module scores compared to those with low *FOXA1*/*SPDEF* (p-value = 7.45 × 10^-5^; Fig. 3J–L). Collectively, these results from integrated scRNA-seq and spatial transcriptomics analyses strongly support that FOXA1 and SPDEF suppress EMT and preserve luminal lineage differentiation in HER2-positive breast cancer cells.

### FOXA1 and SPDEF downregulate the TEAD-YAP-TAZ signaling pathway in HER2-positive breast cancer

To explore how FOXA1/SPDEF loss triggers EMT, we looked for upstream regulators of the EMT gene program. Motif discovery on ATAC-seq peaks that became more accessible upon *FOXA1* or *SPDEF* knockdown yielded an interesting pattern: GATA transcription factor motifs were highly enriched, along with a lesser enrichment of TEA domain transcription factor (TEAD) motifs (Fig. 4A and B). By contrast, FOXA1 and related FOX family motifs, including FOXM1 and FOXK1, were the top hits in regions that lost accessibility in SKBR3-FKD and SKBR3-SPKD cells, as expected since we removed a pioneer factor (*SI Appendix*, Fig. S4A and B). TEAD family transcription factors (TEAD1-4) are effectors of the Hippo signaling pathway and are master regulators of proliferation, development, and cell fate (35–37). We observed increased expression of multiple *TEAD* genes in these knockdown models (Fig. 4C; *SI Appendix*, Fig. S4C and D), and the ATAC-seq analysis confirmed a strong enrichment of TEAD-binding sites in regions of increased chromatin accessibility (Fig. 4D). Thus, loss of FOXA1/SPDEF appears to unleash TEAD activity.

**Figure 4.**
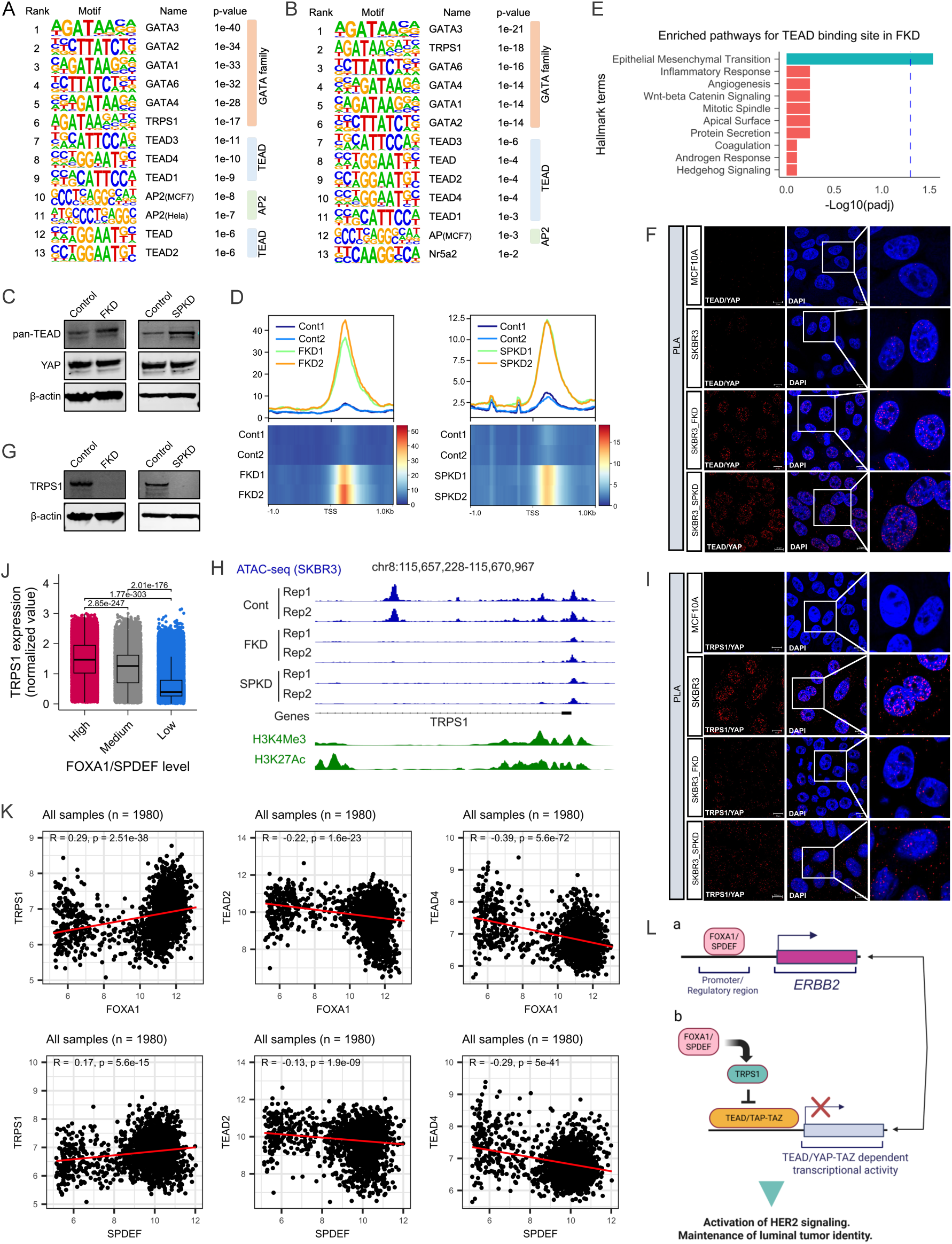
FOXA1 and SPDEF downregulate the TEAD-YAP-TAZ signaling pathway in HER2-positive breast cancer. **A-B**. Top enriched TF binding motifs identified by HOMER known motif analysis associated with up-regulated peak regions from ATAC-seq of SKBR3-FKD (A) and SKBR3-SPKD (B). **C**. Western blots showing the indicated protein levels in SKBR3-FKD and SKBR3-SPKD cells. **D**. Tornado plot illustrating the difference in chromatin accessibility peaks overlapping TEAD family transcription factor binding sites, as identified by ATAC-seq analysis, between control and knockdown conditions. **E**. Bat plot showing the Hallmark ORA results for 245 genes that enriched in TEAD family binding motif in SKBR3-FKD cells compared to SKBR3. **F**. PLA assay of TEAD/YAP in MCF10A, SKBR3, SKBR3-FKD, and SKBR3-SPKD cells. Nuclei stained with DAPI (blue). Scale bars are 10μm. **G**. Western blots showing the TRPS1 protein levels in SKBR3-FKD and SKBR3-SPKD cells. **H**. Genome tracks of *TRPS1* gene promoter region showing the difference in accessibility between control and knockdown cells. Genome tracks showing the identified peaks over pre-defined promoter region (H3K4Me3) and regulatory region (H3K27Ac). **I**. PLA assay of TRPS1/YAP in MCF10A, SKBR3, SKBR3-FKD and SKBR3-SPKD cells. Nuclei stained with DAPI (blue). Scale bars are 10μm. **J**. Boxplot showing the *TRPS1* expression level in *FOXA1*/*SPDEF*^high^ and *FOXA1*/*SPDEF*^low^ expression groups from scRNA-seq data. The median value is indicated by the center line within each box. The lower and upper box edges represent the first and third quartiles, respectively. **K**. Scatter plots illustrating the correlation of gene expression levels between *FOXA1* and *TRPS1*, *TEAD2*, *TEAD4* in the upper three panels, and between *SPDEF* and *TRPS1*, *TEAD2*, *TEAD4* in the lower three panels. Gene expression data were obtained from the METABRIC breast cancer patient cohort. **L**. Schematic model illustrating the cooperative regulatory network involving FOXA1 and SPDEF in *ERBB2* expression and in luminal characteristics in HER2-positive breast cancer. Image was generated by Biorender.

To directly connect TEAD binding to transcriptional outcomes, we analyzed genes associated with newly accessible regions containing TEAD motifs. In SKBR3-FKD cells, 245 genes were identified under such regions and functional analysis of these genes revealed significant enrichment for the Hallmark EMT gene set (adjusted p-value = 1.2 × 10^-2^; Fig. 4E). While a similar trend in SKBR3-SPKD cells did not reach statistically significance (adjusted p-value = 2.06 × 10-1; *SI Appendix*, Fig. S4E), the data collectively link TEAD activation to the EMT gene signature. This aligns with previous studies demonstrating that YAP/TAZ co-activators, when partnered with TEADs, drive pro-EMT and metastatic transcriptional programs (38, 39).

To validate that TEAD-YAP signaling is functionally altered, we performed assays for TEAD and YAP activity. Immunofluorescence analysis showed that TEAD proteins had increased nuclear localization relative to control in SKBR3-FKD and SKBR3-SPKD cells, whereas YAP’s localization did not visibly change (*SI Appendix*, Fig. S4F). Importantly, PLA revealed that TEAD-YAP interactions were significantly increased in both SKBR3-FKD and SKBR3-SPKD cells compared to control cells (Fig. 4F). These findings suggest that even if YAP protein levels remain stable, there is more active TEAD–YAP complex formation when FOXA1/SPDEF are absent, consistent with elevated TEAD transcriptional output driving EMT.

Concurrent with increased TEAD activity in SKBR3-FKD and SKBR3-SPKD cells, we also observed enrichment of TRPS1 binding motifs in accessible chromatin regions. TRPS1, a GATA-type transcriptional repressor commonly found in breast cancers (40), has been known to inhibit YAP/TEAD-dependent transcriptional activity (41–43). ATAC-seq data showed that FOXA1/SPDEF loss reduced chromatin accessibility at the *TRPS1* (Fig. 4H), and robustly decreased TRPS1 expression at the mRNA and protein levels (Fig. 4G, *SI Appendix*, Fig. S4C, D and G). Functionally, this loss diminished YAP-TRPS1 interactions, as shown by PLA, effectively relieving the repression on TEAD (Fig. 4I). The clinical relevance of this axis was supported by scRNA-seq data, where *TRPS1* expression was significantly elevated in *FOXA1*/*SPDEF*^high^ cell populations, compared to *FOXA1*/*SPDEF*^low^ from patient tumors (Wilcoxon rank-sum test, p-value = 1.77 × 10^-303^; Fig. 4J). However, this association was not recapitulated at the spatial transcriptomic level using Visium, which showed no significant correlation between *FOXA1*/*SPDEF* and *TEAD* or *TRPS1* expression across the tumor region (*SI Appendix*, Fig. S4I), suggesting limitations in spatial resolution or context-dependent regulation. Finally, analysis of the METABRIC cohort confirmed a significant positive correlation between *TRPS1* expression and both *FOXA1* and *SPDEF* (R = 0.29 (Pearson), p = 2.51 × 10^-38^ (Pearson) in FKD; R = 0.17 (Pearson), p = 5.6 × 10^-15^ in SPKD). Conversely, expressions of *TEAD2* and *TEAD4* were inversely correlated with the *FOXA1*/*SPDEF* axis (R = -0.22 (Pearson), p = 1.6 × 10^-23^ for *TEAD2* and R = -0.39 (Pearson), p = 5.6 × 10^-72^ for *TEAD4* in FKD; R = -0.13 (Pearson), p = 1.9 × 10^-9^ for *TEAD2* and R = -0.29 (Pearson), p = 5 × 10^-41^ for *TEAD4* in SPKD; Fig. 4K). This clinical data aligns with our mechanistic model, wherein FOXA1/SPDEF promote TRPS1 expression to restrain TEAD activity. Taken together, our findings establish a pathway in which FOXA1 and SPDEF maintain a luminal phenotype by facilitating TRPS1-mediated repression of the TEAD/YAP transcriptional program. The loss of this regulatory axis unleashes TEAD–YAP signaling, driving an EMT program that may enable the transition of luminal tumors to a more invasive state.

## Discussion

FOXA1 is recognized for its ability to regulate the chromatin accessibility of various transcriptional factors, notably the androgen receptor (AR) and ERα (4,44). This realization prompted extensive research into its roles in prostate cancer and ER-positive breast cancer. Although most studies of FOXA1 in breast cancer have focused on its regulation of ER, it is also noted that FOXA1 is expressed in HER2-positive breast cancers with low ERα levels (9). SPDEF is one of the transcriptional factors downstream of FOXA1 (3, 7, 14–16). FOXA1 acts upstream by opening chromatin and enabling SPDEF’s recruitment to target loci, while SPDEF refines lineage-specific transcriptional programs. In many epithelial cells, including breast and prostate, SPDEF promotes luminal differentiation and has been reported to have either tumor suppressive or promotive activity depending on the context (3, 11, 15, 17, 45). This present study provides a more comprehensive understanding of FOXA1 and SPDEF in HER2-positive breast cancer. Interestingly, we found a strong codependency between FOXA1 and SPDEF in HER2-positive tumors. Epigenetic analyses utilizing ChIP-seq and ATAC-seq data indicated that FOXA1 and SPDEF both engage the promoter and an enhancer region of the *ERBB2* gene in HER2-positive SKBR3 breast cancer cells, sites where they do not bind in ER-positive MCF7 breast cancer cells. Knockdown of FOXA1 and SPDEF in SKBR3 cells resulted in diminished *ERBB2* mRNA and reduced HER2 (and EGFR) activation, demonstrating that binding of FOXA1 and SPDEF to the *ERBB2* regulatory sequences is required for full *ERBB2* expression in HER2-positive breast cancer cells. Furthermore, ATAC-seq and RNA-seq analyses revealed that *FOXA1*/*SPDEF* depletion in HER2-positive breast cancer cells decreased *TRPS1* expression, augmented TEAD/YAP interactions, and enhanced the expression of EMT-related genes. Our results thus highlight the crucial role of FOXA1 and SPDEF in sustaining *ERBB2* expression and enforcing luminal tumor identity in HER2-positive breast cancer.

FOXA1 and SPDEF can physically interact, but their transcriptional roles are not identical. They appear to function in a complementary manner within the same regulatory network. FOXA1 is widely recognized as a pioneer factor, capable of binding to compact chromatin and opening regulatory regions to establish a permissive luminal transcriptional framework (4, 10, 46). Within this FOXA1-primed chromatin landscape, SPDEF is recruited to a subset of sites, where it fine-tunes gene expression by activating more specialized programs related to differentiation, secretion, and other context-dependent cellular responses. This can explain why many FOXA1 and SPDEF target genes overlap but are not entirely identical. In essence, FOXA1 ensures broad accessibility of luminal genes and maintains the overall epithelial identity, while SPDEF adds specificity and precision to the transcriptional output. Taken together, these the FOXA1–SPDEF axis functions as a modular regulatory system in which FOXA1 provides the foundational chromatin context and SPDEF refines lineage-restricted pathways. This model accounts for their cooperative interaction yet distinct regulatory footprints, highlighting how multiple layers of transcriptional control reinforce luminal epithelial phenotype.

In breast cancers, FOXA1’s role in facilitating ERα binding to chromatin is well established. It is required for expression of ERα itself and typical estrogen-responsive genes that define the luminal A subtype. Our ChIP-seq and ATAC-seq data in MCF7 cells are consistent with this model, FOXA1 binds to open chromatin at the *ESR1* promoter (*SI Appendix*, Fig. S2E) and presumably elsewhere in the ER network. However, in HER2-positive, ER-negative cells, FOXA1 does not bind to the *ESR1* promoter. Instead, it binds to open chromatin in the *ERBB2* gene. This target switch is emblematic of how FOXA1’s pioneering activity is repurposed in different oncogenic contexts. More broadly, our ChIP-seq demonstrate that many FOXA1 binding sites are exclusive to either MCF7 or SKBR3, while some are shared, indicating FOXA1’s cistrome is context-dependent. The knockdown of SPDEF also led to reduced chromatin accessibility at *ERBB2*, suggesting SPDEF helps maintain *ERBB2* chromatin in an open state. Although FOXA1 and SPDEF both regulate key differentiation oncogenes genes (*ESR1* vs. *ERBB2*), the mechanisms deciding which binding sites FOXA1 uses in each context remain to be identified. It could involve different collaborating factors present in ER-positive versus HER2-positive cells.

Despite FOXA1 being expressed in HER2-positive breast cancers, its molecular interaction with HER2 signaling had not been extensively explored prior to our work (47, 48). Likewise, SPDEF is upregulated in HER2-positive breast cancers compared to normal tissue, but unlike in ER-positive luminal subtypes, its expression in HER2-positive tumors did not clearly correlate with tumor progression or outcome (49). Several studies in other contexts have implicated both FOXA1 and SPDEF in EMT suppression (11, 50–53). Our investigation ties that these observations together by revealing that the FOXA1/SPDEF axis plays a crucial role in maintaining luminal lineage characteristics in HER2-positive breast cancer specifically by repressing chromatin accessibility and the expression of EMT-related genes (37, 39, 54).

A key mechanistic insight from our study is the identification of the TRPS1-TEAD/YAP pathway as a downstream effector of the FOXA1/SPDEF axis. Our data suggest that FOXA1 and SPDEF promote TRPS1 expression, which, in turn, suppresses EMT by antagonizing the activity of TEAD/YAP. However, this finding must be reconciled with previous research that has reported a reciprocal regulatory relationship. For instance, Huang *et al.* found that TRPS1 can induce *FOXA1* expression by binding its promoter in cancer cells (55). Rather than being contradictory, we interpret these results as evidence of a mutual positive feedback loop. In this more sophisticated model, FOXA1/SPDEF and TRPS1 mutually reinforce each other’s expression, creating a robust and self-sustaining transcriptional circuit that locks in the luminal state and actively suppresses cellular plasticity. Disrupting any component of this loop (FOXA1, SPDEF, or TRPS1) leads to a collapse of the entire circuit, consistent with the profound phenotypic shifts to a basal/EMT state that we observe upon depletion of FOXA1 and SPDEF. This feedback loop concept aligns with the idea of lineage surveillance. As long as FOXA1/SPDEF and TRPS1 keep each other active, the cell is confined to a luminal epithelial identity but once the loop is broken, the cell can transit to a different state (with TEAD/YAP taking over, driving mesenchymal programs).

Motif analysis of ATAC-seq changes revealed an interesting predominance of GATA factor in regions that opened upon FOXA1/SPDEF loss. GATA3 is another luminal-lineage TF often co-expressed with FOXA1 in ER-positive breast cancer. GATA3 and FOXA1 co-bind many sites in ER-positive cells to maintain luminal chromatin (3, 56, 57). In the context of HER2-positive tumors, GATA3 expression is low, but other GATA factors, such as TRPS1, come into play. It’s worth noting that TRPS1, despite being a “GATA” acts mainly as a repressor. The fact that GATA motifs appeared in increased-accessibility regions in our data suggests that when FOXA1/SPDEF are gone, some GATA-binding factors might be losing binding. This is somewhat speculative, but could be discussed as an avenue. How exactly the network of luminal TFs (FOXA1, GATA3, TRPS1, etc.) coordinates to maintain identity is complex and perhaps involves collaborative repression and activation.

Luminal identity in breast cancer is maintained by a network of lineage-defining transcription factors, such as FOXA1, GATA3, and SPDEF, that sustain open chromatin at luminal enhancers (3, 56, 57). These factors directly support the expression of luminal oncogenes, including *ESR1* and *ERBB2*, thereby coupling the differentiation state with oncogenic signaling (57, 58). In ER-positive tumors, FOXA1 and GATA3 facilitate ERα’s recruitment to chromatin, reinforcing estrogen-driven transcriptional programs. In HER2-positive tumors, now we see that luminal factors like FOXA1/SPDEF stabilize HER2-driven signaling and preserve epithelial differentiation. Loss of these luminal regulators leads to diminished *ESR1*/*ERBB2* expression and a shift toward basal or stem-like states, highlighting the interdependence between luminal identity and luminal oncogene activity.

This study dissects a FOXA1/SPDEF circuit primarily in SKBR3 cells; validating generality across diverse HER2-positive models and *in vivo* will strengthen external validity. The proposed FOXA1/SPDEF–TRPS1 positive feedback loop and the direct regulation of *ERBB2* are supported by accessibility, FOXA1 ChIP-seq, and expression changes, yet would benefit from targeted perturbations (CRISPRi/a of the *ERBB2* regulatory elements we map, enhancer–reporter assays, and genetic rescue of TRPS1) to close remaining causal gaps. While TEAD/YAP activation gains support, further studies are needed to clarify the sufficiency and necessity of TEAD/YAP activation, increased TEAD levels, and TEAD– YAP PLA. Genome-wide occupancy profiling (of TEADs, YAP/TAZ, and TRPS1) and pharmacologic/genetic TEAD–YAP inhibition experiments will help elucidate these aspects. Our spatial and single-cell analyses are constrained by Visium spot resolution and the use of imputation; cross-platform replication mitigates, but does not eliminate, analytical bias. Finally, pooled shRNA knockdowns can carry off-target effects; orthogonal CRISPR-based approaches, acute degron systems, functional EMT phenotyping (invasion/migration, plasticity assays), and therapeutic response tests (e.g., anti-HER2 sensitivity upon circuit disruption) are priorities to translate this axis into biomarkers or targets.

In conclusion, our study demonstrates that FOXA1 and SPDEF play a crucial role in preserving luminal characteristics, reducing EMT, and enhancing *ERBB2* expression in HER2-positive breast cancer cells. We have defined a novel, ER-independent regulatory network wherein a FOXA1/SPDEF axis, stabilized by a positive feedback loop with TRPS1, acts as a master guardian of luminal identity. This axis simultaneously drives expression of the core oncogene *ERBB2* while actively suppressing a latent TEAD/YAP-driven mesenchymal program (Fig. 4L). These findings are consistent with prior reports demonstrating that FOXA1/SPDEF expression helps distinguish luminal versus basal subtypes (59, 60). A deeper understanding of the molecular interactions between FOXA1, SPDEF, and TRPS1 in HER2-positive tumors may pave the way for the development of novel therapeutic strategies for this tumor subtype. For example, interventions that reinforce this luminal circuit to prevent EMT and resistance or, conversely, therapies that target the YAP/TEAD pathway could be explored. Overall, our work sheds light on the luminal-basal plasticity in breast cancer and identifies the FOXA1–SPDEF–TRPS1 axis as a potential vulnerability that might be leveraged to improve outcomes in HER2-positive breast cancer.

## Materials and Methods

### Cell culture

The human cell lines, SKBR3 and MCF7, were obtained from ATCC and maintained in culture in DMEM +GlutaMAX-1 (Gibco-Life Technologies) containing 10% fetal bovine serum (FBS) and pen/strep (Gibco-Life Technologies) at 37°C in 5% CO_2_. MCF10A cells were cultured in DMEM/F12 (Gibco-Life Technologies) containing 5% horse serum, EGF (100ng/ml), hydrocortisone (1mg/ml), cholera toxin (1mg/ml), insulin (10μg/ml), and pen/strep (Gibco-Life Technologies) at 37°C in 5% CO_2_.

### Knockdown Cell Line

A stable cell line expressing shRNA directed against FOXA1 and SPDEF was generated by transducing cells with commercially prepared lentiviruses containing three individual shRNA directed against FOXA1 mRNA (sc-37930-V) (Santa Cruz) and SPDEF mRNA (sc-s5845-V). Cells were cultured in 6-well plates and infected by adding the shRNA lentiviral particles to the culture for 48 hours per the manufacturer’s instructions. Stable clones expressing the specific shRNAs were selected using 5μg/ml of puromycin (Gibco-life technologies) and pooled to generate the cells used in the experiments.

### Immunofluorescence

Cells were grown on coverslips, fixed in 4% paraformaldehyde for 20 min, permeabilized with 0.2 % Triton X100 for 10 mins, washed three times with PBS, and incubated with primary antibody overnight at 4°C. The cells were washed three times with PBS and incubated with a secondary antibody for 1 hour at room temperature. After washing, coverslips were mounted using Prolong Gold antifade reagent with DAPI (Invitrogen).

Confocal microscopy images were obtained using a Zeiss LSM 880 confocal microscope. Primary antibodies included those against FOXA1 (ab170933) from Abcam (Waltham, MA); SPDEF from NOVUS Biologicals (Centennial, CO); HER2 (MA1-35720) and TRPS1 (PA5-84874) from Thermo Scientific (Waltham, MA); phosphor-HER2 and pan-TEAD (13295) from Cell Signaling (Danvers, MA); YAP (sc-101199) from Santa Cruz (Dallas, TX).

### RNA Extraction and Real-Time RT-PCR

RNA was isolated using TRIzol (Invitrogen). Quantitative RT-PCR was performed with the SuperScript III Platinum One-Step qRT-PCR Kit (Invitrogen) using a Step One Plus Real-Time PCR System (Applied Biosystems) and the following TaqMan primer sets: FOXA1, Hs04187555_m1; SPDEF, Hs0017942_m1; ErbB2, Hs01001580_m1. Human HPRT1 (4326321E) (Invitrogen) was used as a reference gene. Relative mRNA expression was determined using Step One Software v2.2.2 (Applied Biosystems).

### Immunoblotting

Protein samples were prepared from cells using standard methods, subjected to SDS-PAGE, and transferred to a nitrocellulose membrane by wet western blot transfer (Bio-Rad). The membrane was blocked in TBST buffer (TBS + 1% Tween) containing 5% milk for 1 hour at room temperature. The blocked membranes were incubated overnight at 4°C with specific primary antibodies (Odyssey blocking buffer, 927-40000). The membranes were washed three times with TBST buffer and then incubated with specific secondary antibodies provided by LI-COR for two hours at room temperature. After three washes with TBST buffer, the membranes were analyzed using the ODYSSEY Infrared Imaging system (LI-COR). All immunoblot experiments were performed at least three times, and representative blots are shown in the figures. Primary antibodies included those against FOXA1 (ab170933) from Abcam (Waltham, MA); SPDEF from NOVUS Biologicals (Centennial, CO ); phospho-HER2 (thr1221/1222) (2243S), EGFR (4267S), and pan-TEAD (13295) from Cell Signaling (Danvers, MA) from Cell Signaling (Danvers, MA); HER2 (sc-33684), phospho-EGFR (sc-12351) from Santa Cruz (Dallas, Texas); TRPS1 (PA5-84874) from Thermo Scientific (Waltham, MA).

### Single cell RNA sequencing data analysis

Count matrices from published single cell RNA sequencing (scRNA-seq) datasets (25) were downloaded from the NCBI Gene Expression Omnibus (GSE161529) and then analyzed using Seurat version 4.0 as described (61). 15 ER-positive and 6 HER2-positive patient-derived datasets were included for analysis. Cells were divided by a high group and low group, depending on their MAGIC imputation counts (t = 2) of *FOXA1*, *SPDEF*, *ESR1* and *ERBB2* expression (62). The high group was assigned by having an expression level greater than median, and remaining cells were designated as the low group, unless otherwise noted. P-value was calculated from a down-sampled dataset by 1,000 to 2,000 cells to avoid zero inflation of scRNA-seq data.

### Spatial transcriptomics data analysis

The Visium and Xenium dataset used in this analysis were provided by 10X Genomics. Human breast cancer sections are freely available datasets and can be downloaded at from the 10X website (https://www.10xgenomics.com/resources/datasets). Visium spots were annotated refer to the deconvolution method Robust Cell Type Decomposition (RCTD) and previous report (26). To calculate the module score for EMT and TEAD feature expression program per spot, Seurat’s AddModuleScore function was applied to Visium data. For EMT module score, 136 genes were selected because they were both involved in Hallmark EMT pathway gene set and have comparable count read (63).

### ChIP-seq and data analysis

ChIP-seq for FOXA1 was performed as described in (64, 65). (FOXA1 antibody; cat. ab170933). DNA was purified with a QIAquick PCR purification kit (Qiagen, Germantown, MD, USA) for sequencing. ChIP-seq library preparation was done using the TruSeq Nano DNA Library Prep Kit (Illumina, San Diego, CA, USA) and sequenced on the Illumina NextSeq550 sequencing platform (performed at the Yale Sequencing Center). The quality of raw sequencing reads was evaluated with FastQC (https://www.bioinformatics.babraham.ac.uk/projects/fastqc/). The raw sequencing reads from FOXA1 and matched input DNA were mapped to the human reference genome (hg38) using the Burrows-Wheeler Aligner (BWA) mem program (version 0.7.9a) (66). After alignment, PCR duplicates were removed with Picard tools (http://broadinstitute.github.io/picard/) (MarkDuplicates v.2.9.0) and FOXA1 binding regions (i.e., peaks) were identified using MACS tool (version 1.4). Peaks were annotated using annotatePeaks.pl script from HOMER. Deeptools (version v.3.1) and Integrative Genomics Viewer (IGV, version 2.12) were used for visualization.

### ATAC-seq and data analysis

ATAC-seq was performed as described in ref (67). Extracted cells from SKBR3 and SKBR3-FKD with two biological replicates were subsequently processed using standard DNA extraction and library preparation protocols (performed at the Yale Sequencing Center). Libraries were prepared according to the recommended Illumina NextSeq550 sequencing platform (performed at the Yale Sequencing Center).

ATAC-seq data processing was achieved as previously described (68). HOMER software was used for motif analysis based on the peak annotation files. The findMotifsGenome.pl script in HOMER was used to discover enriched motifs for each condition. To examine the chromatin accessibility changes around the peak summit, deepTools (version 3.5) was used. Heat map (tornado plots) of peaks were generated by combining scores for regions and binning the region from -1 kb to +1 kb of transcription start site. Differential accessibility of the peaks was assessed using DESeq2 (version 3.16) using peak scores between pairwise comparisons of SKBR3-FKD to SKBR3.

### Bulk RNA-seq and data analysis

The raw sequencing reads files in Fastq format were assessed for quality with FastQC. Pseudo-alignment was achieved using Salmon (version 1.9) with hg38 reference genome. Subsequent analyses were performed using DESeq2 (version 3.16) with output quantification files. The differential expression of SKBR3-FKD and SKBR3-SPKD samples was performed using DESeq2 with R package. Adjusted P-value of 0.05 by Benjamini-Hochberg method and log2FC value of 1 are considered as significant, unless otherwise noted. Functional pathway enrichment of differentially expressed genes were determined using clusterProfiler package (version 4.4) from the molecular signature database Hallmark collection and Oncogenic signature (version 7.4).

### Genetic dependency analysis

The dependency data is available online, at https://depmap.org/portal/. RNAi and CRISPR screening data for 1,255 cell lines were used to identify cellular viability following gene perturbation. Pearson correlation between each gene’s log2(TPM+1) expressions from 22Q2 Public were computed. Genes were examined to compare distinct dependency pattern according to lineage sub-subtype in breast cancer.

### Proximity Ligation Assays

Proximity ligation assays (PLA) were performed using the Duolink™ assay kit (Sigma). SKBR3 and MCF10A cells were seeded onto 22 mm round collagen-coated coverslips (Corning, Cat N 354089). The experiments were performed when the cells reached 80% confluence. The cells were washed 3 times with PBS and paraformaldehyde (4% in PBS) was added to each well for 20 min, after which they were permeabilized with 0.2% Triton X100 in PBS. Permeabilized cells were incubated with combinations of the following antibodies: FOXA1 (ab170933) from Abcam (Waltham, MA); SPDEF from NOVUS Biologicals (Centennial, CO); TRPS1 (PA5-84874) from Thermo Scientific (Waltham, MA), pan-TEAD (13295) from Cell Signaling (Danvers, MA) and YAP (sc-101199) from Santa Cruz (Dallas, TX). PLA probes were then added and the assay was performed as per the manufacturer’s instructions. All images were obtained using a Zeiss LSM 880.

### Quantification and statistical analysis

The number of repeats (n) refers to the number of independent experimental observations in the legend of each figure. Statistical analyses were performed with Prism 6.0 (GraphPad Software, La Jolla, CA). Statistical significance was determined using an unpaired t test for comparisons between two groups. Group data were presented as mean±SEM. The P-values less than 0.05 (indicated by *), or less than 0.01 (**), or less than 0.001 (***), or less than 0.0001 (****) were considered as significant.

## Data availability

ChIP-seq, RNA-seq, and ATAC-seq data from FOXA1 knockdown experiments in HER2-positive SKBR3 cells have been deposited in the NCBI Gene Expression Omnibus (GEO) under accession numbers GSE267919, GSE267920, and GSE267921, respectively. RNA-seq and ATAC-seq data from SPDEF knockdown experiments are prepared for deposition in GEO; however, submission is currently delayed due to the US government shutdown affecting NCBI services. These datasets will be deposited immediately upon resumption of GEO operations, and accession numbers will be provided before publication.

## Author Contributions

Study concept and design: J.Jeong, J. Wysolmerski and J.Choi.; acquisition of data: J.Jeong, J.Lee, and J.Choi.; analysis and interpretation of the data: J.Jeong, J.Lee, K.Yoo, S.Kim, J.Lim and J.Choi. technical or other material support: J.Shin, J.Kim, Y.Tanaka, H.S.Tae, and I.Park.; writing and review of the manuscript: J.Jeong, J.Lee, J. Wysolmerski and J.Choi.

## Acknowledgments

This work was partly supported by the National Research Foundation of Korea (NRF) Grant funded by the Korean government (RS-2023-00212238, J.Choi; RS-2024-00461521 to J.Lee). R01 HD100468-01A1 and R01 HD076248-06A1 from the NIH to J. Wysolmerski. JC was supported by the MSIT(Ministry of Science and ICT), Korea, under the ICAN(ICT Challenge and Advanced Network of HRD) program(IITP-2024-RS-2022-00156439) supervised by the IITP(Institute of Information & Communications Technology Planning & Evaluation).

