## Supporting Information for "A FOXA1/SPDEF Co-regulatory Axis Drives ERBB2 and Suppresses TEAD/YAP-Driven EMT to Maintain Luminal Identity in HER2+ Breast Cancer"

Fig. S1. Reciprocal Co-dependency of FOXA1 and SPDEF in HER2-positive breast cancer.

**A**. Jitter plot showing the distribution of FOXA1 gene effect scores (Chronos scores) across various cancer model subtypes from the DepMap dataset. Each group represents a distinct model subtype, highlighting the differential dependency of cell lines on FOXA1. **B**. UMAP plot showing the distribution of ER-positive and HER2-positive cell populations in scRNA-seq data. **C**. UMAP plot displaying the distribution of annotated cell types including immune cells, epithelial cells, and fibroblasts in the scRNA-seq dataset. **D**. Feature plot showing the representative cell type-specific markers. **E**. Spatial distribution of annotated cell types detected in Visium spatial transcriptomics data. **F**. Spatial plots showing a normalized expression of FOXA1, SPDEF, and ErbB2 in the histological image. **G**. Immunofluorescence images of FOXA1 or SPDEF (green) in SKBR3 cells. Nuclei stained with DAPI (blue). Scale bars are 10μm.

Fig. S2. FOXA1 and SPDEF are required for ERBB2 expression in HER2-positive breast cancer cells.

**A-B**. Dot plots showing the Oncogenic signature gene sets significantly enriched (adjusted p-value < 0.05, Benjamini-Hochberg correction) in the SKBR3-FKD cells (A) and SKBR3-SPKD cells (B) compared to the SKBR3. The left panel display upregulated gene sets, while the right panels show downregulated gene sets. **C-D**. Volcano plots showing the differential gene expression profile in SKBR3-FKD (C), and SKBR3-SPKD (D) versus SKBR3 cells. Red and blue dots indicate genes from the ERBB2_UP_V1_UP and ERBB2_UP_V1_DN leading edge subset, respectively. **E**. Genome tracks of *ESR1* gene promoter region showing the difference accessibility between control and knockdown cells. Genome tracks showing the identified peaks over pre-defined promoter region (H3K4Me3) and regulatory region (H3K27Ac). FOXA1 binding motifs matched with ChIP-seq peak regions were marked with a green square.

Fig. S3. FOXA1 and SPDEF are required for luminal identity in HER2-positive breast cancer.

**A-B**. Dot plots showing the Hallmark signature gene sets significantly enriched (adjusted p-value < 0.05, Benjamini-Hochberg correction) in the SKBR3-FKD cells (A) and SKBR3-SPKD cells (B) compared to the SKBR3. The left panel display upregulated gene sets, while the right panels show downregulated gene sets. **C-D**. Dot plots illustrating the significantly enriched Hallmark signature terms of GSEA in the analysis of SKBR3-FKD cells (C) and SKBR3-SPKD cells (D) compared to the SKBR3. **E-F**. Volcano plots showing the differential gene expression profile in SKBR3-FKD (C), and SKBR3-SPKD (D) versus SKBR3 cells. Red dots indicate genes from the HALLMARK_EMT leading edge subset. **G-H**. Tornado plots illustrating the differential chromatin accessibility peaks identified by ATAC-seq between control and knockdown conditions. ATAC-seq reads in 2-kb regions centered on the TSS sites of annotated promoter-TSS sites of protein-coding genes. **I**. Dot plot showing the significantly enriched Hallmark gene sets by ORA performed for DEGs (|Log2FC| > 1.5) in the SKBR3-SPKD cells compared to the SKBR3. Genes corresponding with DARs annotated only as Promoter-TSS were used for analysis. **J**. Dot plot showing the Hallmark signature gene sets significantly enriched (adjusted p-value < 0.05, Benjamini-Hochberg correction) in the SKBR3-FKD cells compared to the SKBR3. The left panel display upregulated gene sets, while the right panels show downregulated gene sets. Genes corresponded with DARs annotated only as Promoter-TSS were used for analysis. **K**. Dot plot illustrating the top 10 enriched terms of GSEA in the analysis of SKBR3-FKD cells compared to the SKBR3. Genes corresponded with DARs annotated only as Promoter-TSS were used for analysis. **L**. Scatter plot showing the correlation of the log2FC values between RNA-seq and ATAC-seq of SKBR3-SPKD. Displayed R value determined by Pearson correlation. **M-N**. Scatter plots showing the correlation between the log2FC values of HALLMARK_EMT genes derived from RNA-seq and ATAC-seq in SKBR3-FKD and SKBR3-SPKD cells. Displayed R value determined by Pearson correlation.

Fig. S4. FOXA1 and SPDEF downregulate the TEAD-YAP-TAZ signaling pathway in HER2-positive breast cancer.

**A-B**. Top enriched TF binding motifs identified by HOMER known motif analysis associated with down-regulated peak regions from ATAC-seq of SKBR3-FKD (A) and SKBR3-SPKD (B). **C-D**. Volcano plots illustrating differential gene expression profiles in SKBR3-FKD (C), and SKBR3-SPKD (D) cells compared to SKBR3 controls. Genes belonging to the *TRPS1* and *TEAD* family are highlighted to emphasize their expression changes. **E**. Bat plot showing the Hallmark ORA results for 115 genes that enriched in TEAD family binding motif in SKBR3-SPKD cells compared to SKBR3. **F-G**. Immunofluorescence images of TEAD, YAP (F) and TRPS1 (G) (red) in MCF10A, SKBR3, SKBR3-FKD and SKBR3-SPKD cells. Nuclei stained with DAPI (blue). Scale bars are 10μm. **H**. Boxplots showing the expression levels of *TEAD* family genes in *FOXA1*/*SPDEF*^high^ and *FOXA1*/*SPDEF*^low^ expression groups from scRNA-seq data. The median value is indicated by the center line within each box. The lower and upper box edges represent the first and third quartiles, respectively. **I**. Spatial plots showing a normalized expression of *TEAD* family genes and *TRPS1* in the histological image.
