## Supplementary figures and images for "A FOXA1/SPDEF Co-regulatory Axis Drives ERBB2 and Suppresses TEAD/YAP-Driven EMT to Maintain Luminal Identity in HER2+ Breast Cancer"

### Fig S1

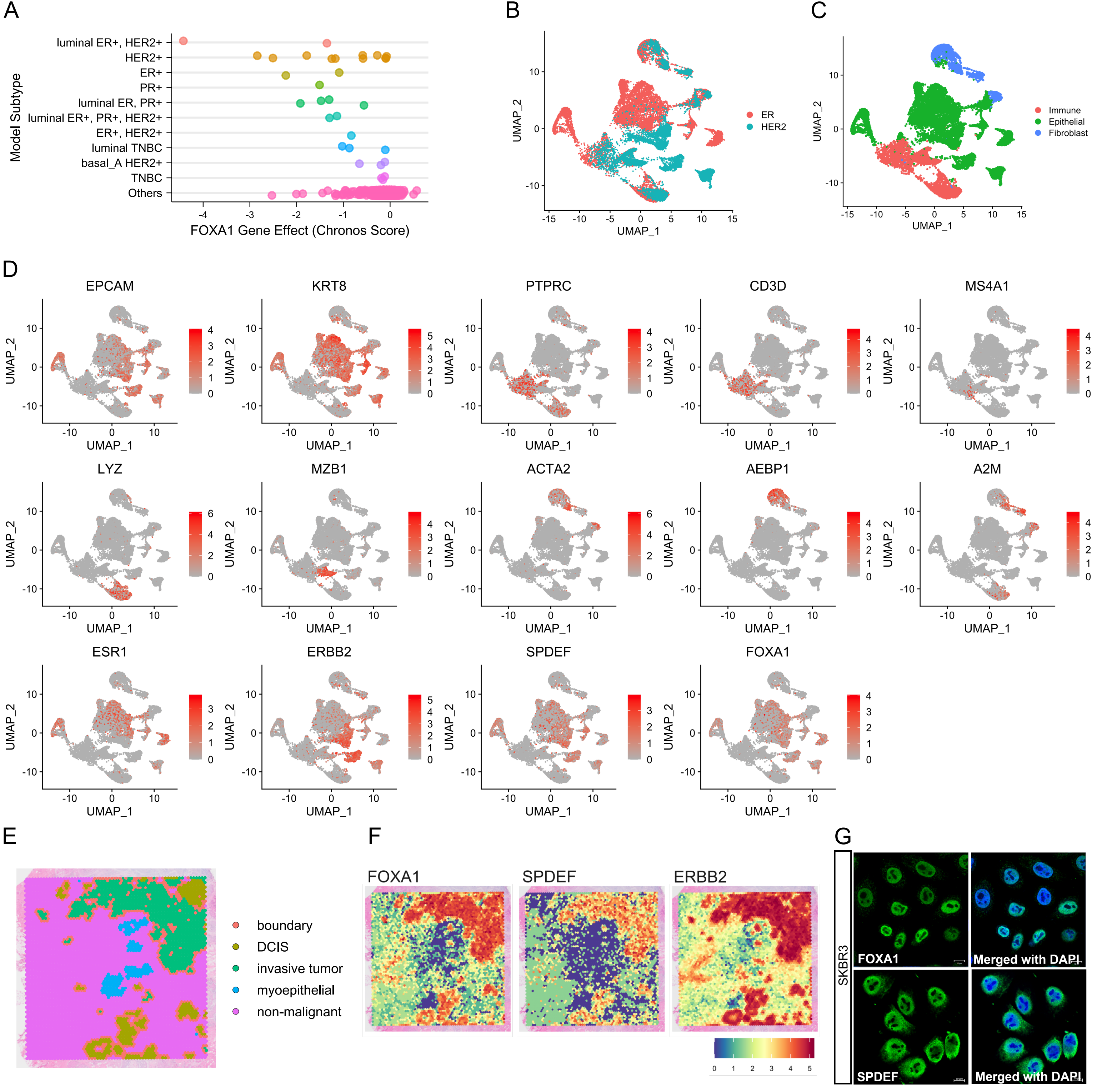

### Fig S2

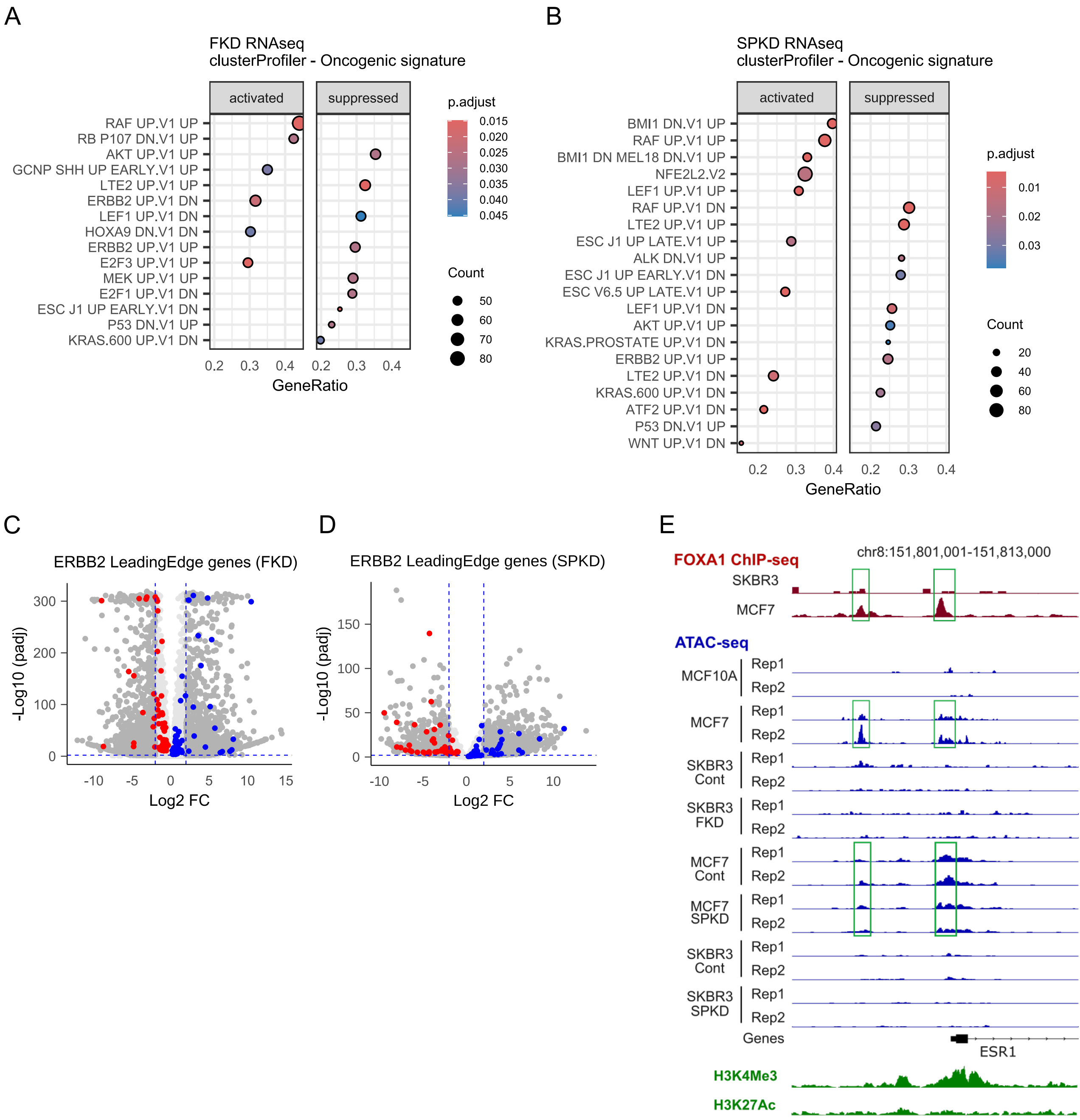

### Fig S3_1

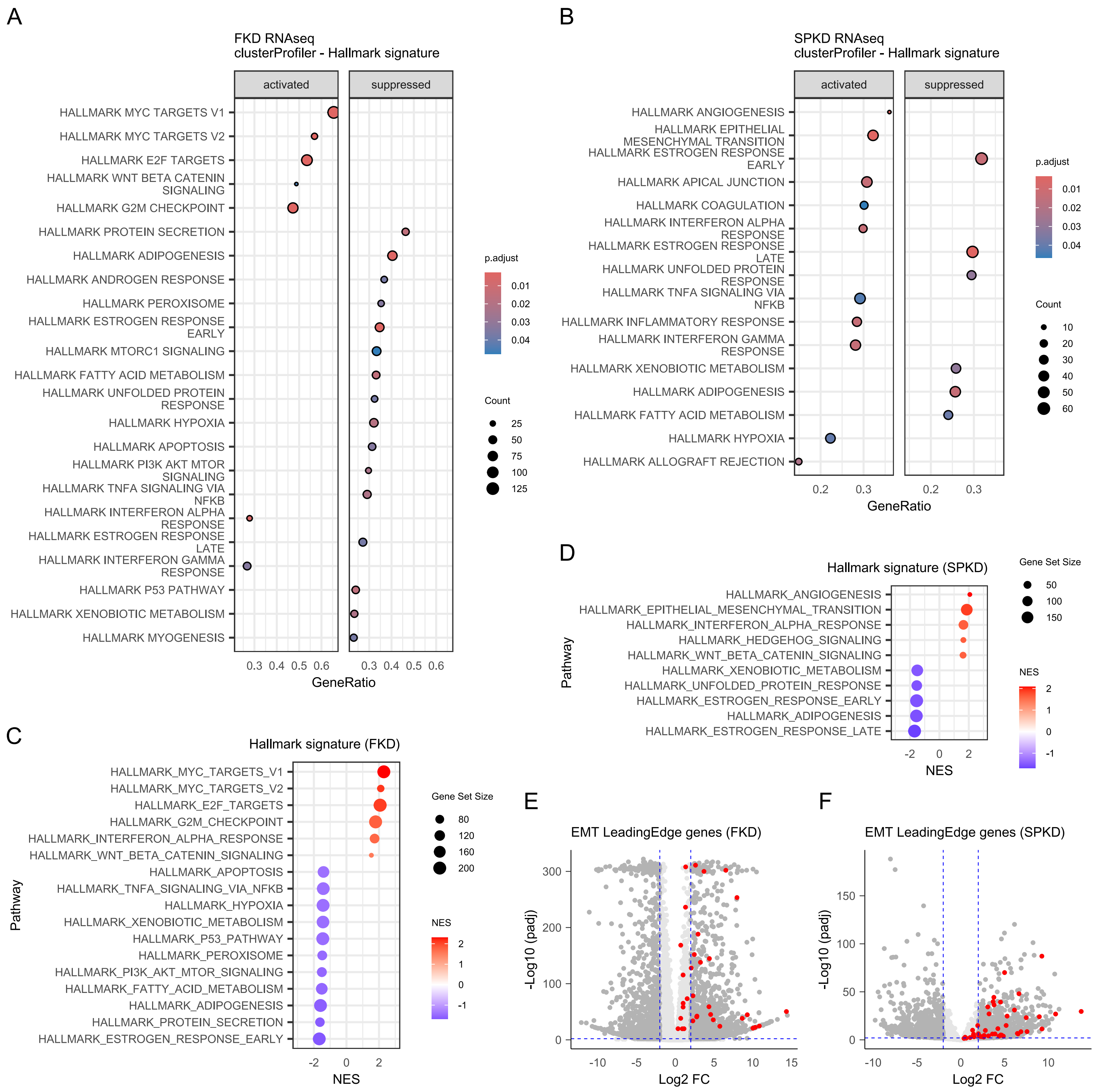

### Fig S3_2

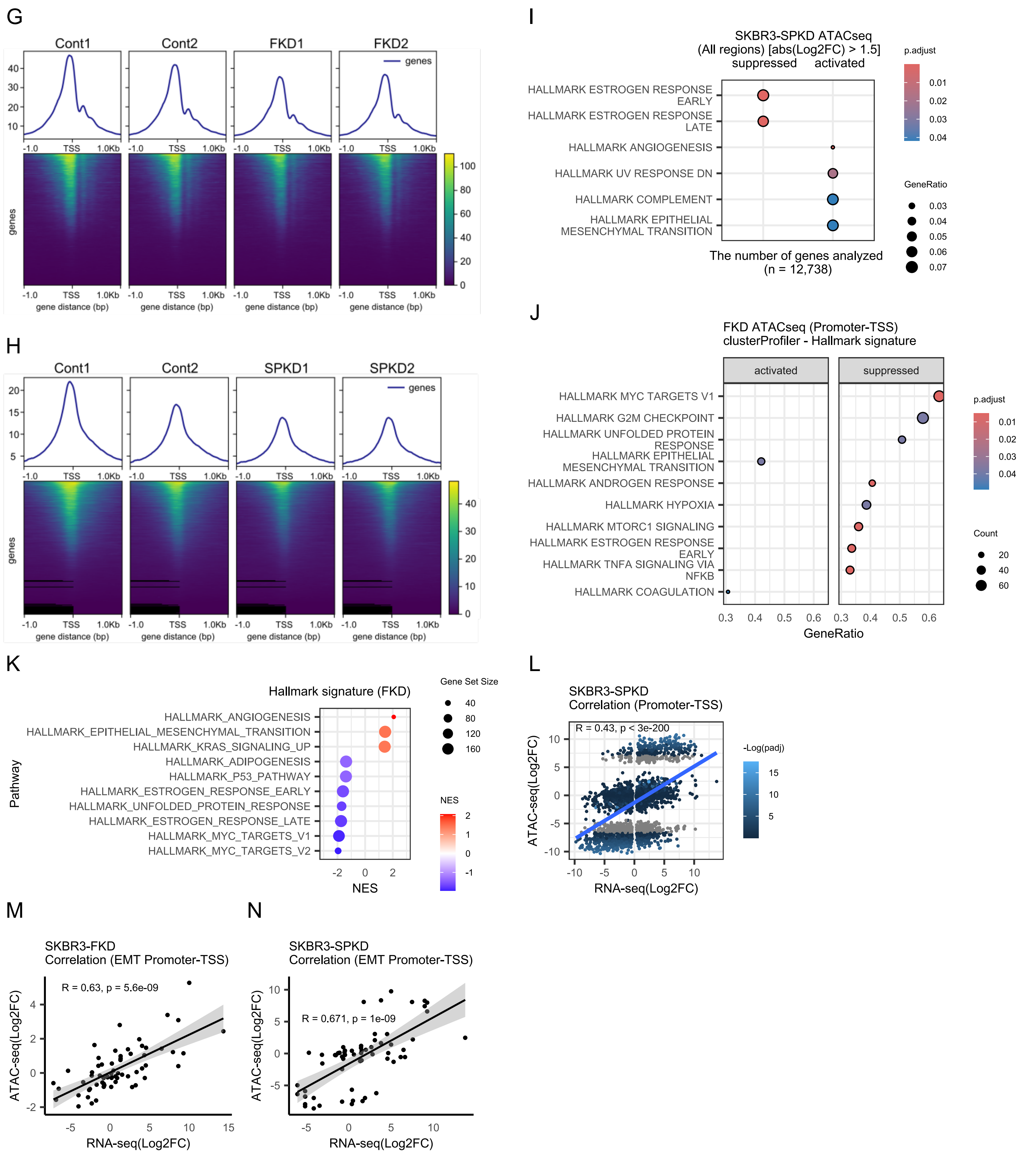

### Fig S4

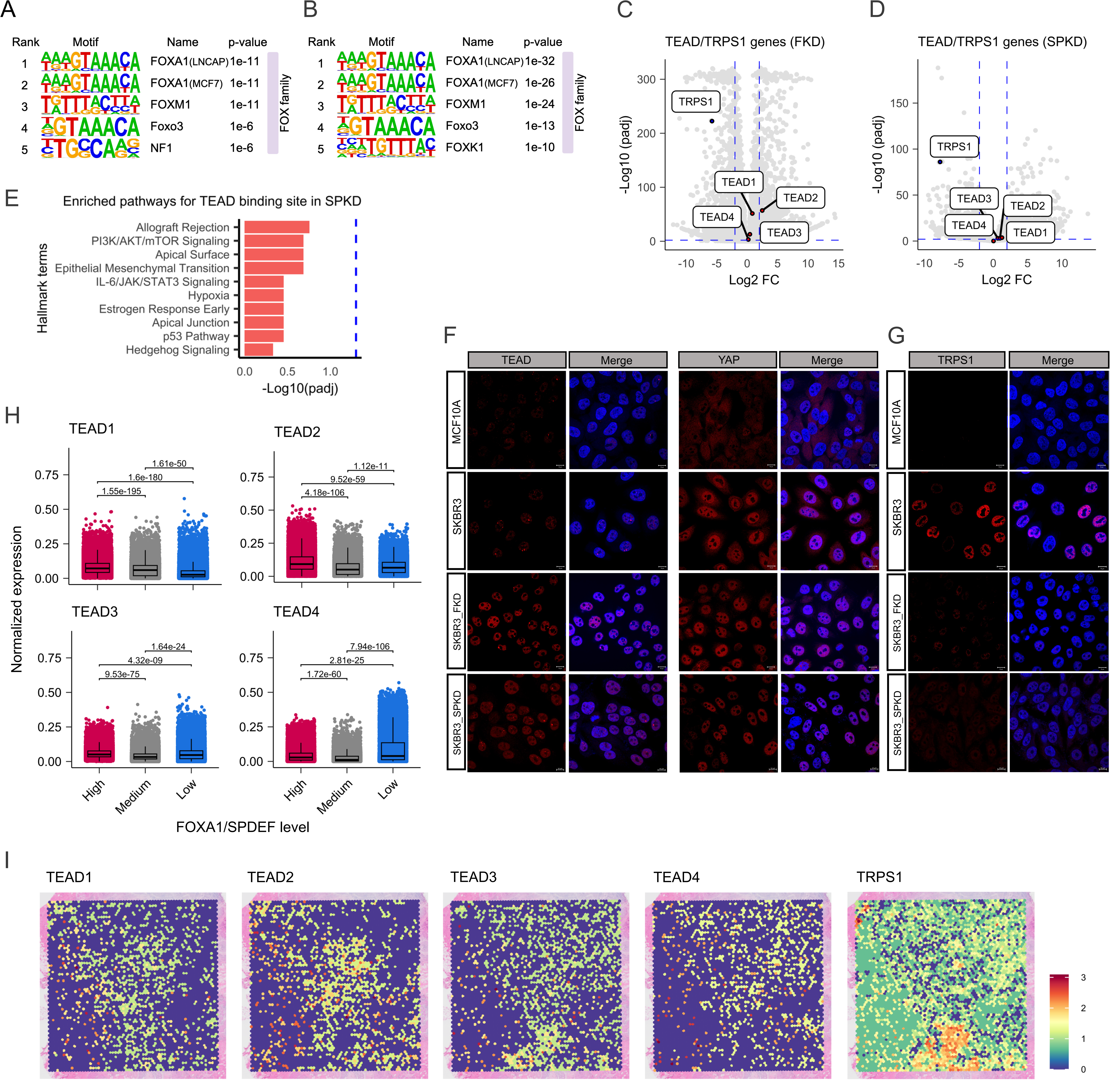
